# Aberrant DEGS1 sphingolipid metabolism impairs central and peripheral nervous system function in humans

**DOI:** 10.1101/347591

**Authors:** Gergely Karsai, Florian Kraft, Natja Haag, G Christoph Korenke, Benjamin Hänisch, Saranya Suriyanarayanan, Regula Steiner, Cordula Knopp, Michael Mull, Markus Bergmann, J Michael Schröder, Joachim Weis, Miriam Elbracht, Matthias Begemann, Thorsten Hornemann, Ingo Kurth

**Author notes:** Correspondence and requests for material should be addressed to I.K. or to T.H. These authors contributed equally to this work (Gergely Karsai, Florian Kraft, Natja Haag, Thorsten Hornemann, Ingo Kurth).

## Abstract

Sphingolipids including ceramides are important components of cellular membranes and functionally associated with fundamental processes such as cell differentiation, neuronal signaling and myelin sheath formation. Defects in the synthesis or degradation of sphingolipids are associated with various neurological pathologies, however, the entire spectrum of disorders affecting sphingolipid metabolism remains elusive. By whole-exome sequencing in a patient with a multisystem neurological disorder of both the central and peripheral nervous system, we identified a homozygous variant p.(Ala280Val) in *DEGS1,* encoding an enzyme of the ceramide synthesis pathway. The blood sphingolipid profile and patient-derived fibroblasts both showed a significant shift from the unsaturated to the dihydro-forms of sphingolipids. Moreover, an atypical and potentially toxic sphingolipid metabolite is formed as consequence of the altered synthesis pathway. The changes in the sphingolipid profile were recapitulated in a CRISPR/Cas-based *DEGS1* knockout HAP1-cell model and by chemical inhibition of DEGS1, suggesting a loss of DEGS1 function in the disease. DEGS1 insufficiency is thus a novel cause for a multisystem neurological disorder. A sphingolipid-rich diet may correct the metabolic profile and improve the clinical outcome of affected individuals and suggests that this heritable condition might be treatable.

**Abbreviations:**

SL: Sphingolipids
SPT: serine-palmitoyltransferase
Cer: Ceramides
dhCer: dihydroceramide
S1P: sphingosine-1-phosphate
SO: sphingosine
HSAN: hereditary sensory and autonomic neuropathy

---

Sphingolipids (SL) are a ubiquitous and structurally diverse class of lipids and potent signaling molecules. Together with sterols and glycerosphospholipids they are major constituents of eukaryotic membranes, and assemble with cholesterol in lipid domains to maintain membrane function. SLs supply essential components of the myelin sheath ^1^ and are thus of fundamental importance for the function of nerves. Whereas in the central nervous system (CNS) oligodendrocytes support and insulate neurons, equivalent function is provided by Schwann cells in the peripheral nervous system (PNS). However, besides a role in myelin formation and maintenance, signaling SLs such as sphingosine-1-phosphate (S1P) and ceramide-1-phosphate are bioactive lipid hormones that regulate a variety of physiological functions including cell migration and viability, telomere stability, angiogenesis, and many others. Ceramides are the central components of the SL metabolism as they form the building blocks for complex SLs like sphingomyelins and glycosylceramides. They also represent the crossroad for the degradation and salvage pathways ^2^. Ceramide biosynthesis starts at the endoplasmic reticulum (ER) with the conjugation of L-serine and palmitoyl-CoA, the rate-limiting step catalyzed by serine-palmitoyltransferase (SPT). The immediate product (3-keto-sphinganine, 3-ketoSA) is reduced to sphinganine (SA) which is then *N*-acylated to dihydroceramide (dhCer) by one of six ceramide synthases (CerS1-6) ^3^. The final step of the de novo pathway is catalyzed by the Δ4-dihydroceramide desaturase (DEGS1) that introduces a Δ4,5-trans (Δ 4,5E) double bond into the sphinganine backbone of dhCer thereby forming ceramide. On the catabolic side, ceramides are deacylated by ceramidases to form sphingosine (SO) ^2 4^, which is vital for the cell as it can be either recycled in the salvage pathway back to ceramides or phosphorylated by sphingosine kinases (SK1/SK2) to form S1P. S1P is a potent lipid hormone that binds to six S1P receptors (SP1R1-6), which are involved in a multitude of cellular responses. Metabolically, S1P can either be converted back to SO through action of S1P phosphatases, (S1PPase), or terminally degraded to trans-2-hexadecenal and ethanolamine phosphate by the S1P lyase (SGPL1) ^5^. To a minor extend sphinganine can also be phosphorylated and degraded by this pathway.

For the formation of complex SLs, ceramides are transported from the ER to the Golgi where they get converted into phosphosphingolipids (sphingomyelins) or glycosphingolipids (gal-Cer, glu-Cer and gangliosides) ^6^. Importantly, complex sphingolipids can be formed from both dhCer and Cer, forming either dihydrosphingolipids (dhSL) with a saturated or typical sphingolipids (SL) with a Δ4,5E sphingoid base. However, dhSL are usually minor and typically make up < 10% of the total SL depending on cell type and tissue.

The degradation of complex SLs requires dedicated catabolic enzymes, such as glycohydrolases and sphingomyelinases that reside in the plasma membrane, ER, Golgi apparatus and lysosomes ^7, 8^.

Defects in these enzymes cause a variety of pathological conditions ^9^. Lipid storage diseases such as M. Fabry, M. Gaucher, M. Farbers, M. Niemann-Pick and M. Tay-Sachs are primarily SL storage disorders due to defects in SL catabolism. Besides defects in the catabolism of complex sphingolipids, defects in the de novo synthesis are also associated with disease. Mutations in *SPTLC1* and *SPTLC2* cause hereditary sensory and autonomic neuropathy type I (HSAN1) which is associated with the formation of an aberrant and neurotoxic class of sphingolipids (1-deoxysphinglipids) (OMIM #162400, #613640). Mutations in ceramide synthase 1 and 2 (*CERS1/2*) were associated with progressive myoclonic epilepsy, generalized ^10, 11, 12^ tonic-clonic seizures, tremor, dysarthria, ataxia and developmental delay ‘ ‘. Mutations in the GM3 synthase gene *ST3GAL5* were associated with refractory epilepsy, myoclonus, generalized tonic-clonic seizures, psychomotor delay, developmental stagnation, blindness, and deafness due to an increase in LacCer and other gangliosides (OMIM #609056) ^13, 14^, whereas defects in the GM2/GD2 synthase (*B4GALNT1*) lead to GM3 accumulation and a complex form of hereditary spastic paraplegia with cognitive impairment and seizures (OMIM #609195) ^15^. Recently, mutations in *SGPL1* were associated with a spectrum of disease phenotypes ^16, 17, 18^ including recessive steroid-resistant nephrotic syndrome (SRNS), ichthyosis, adrenal insufficiency, immunodeficiency and brain defects (OMIM #617575). In addition, *SGPL1* mutations were found in a family with Charcot-Marie-Tooth neuropathy (CMT). The mutation segregated with the disease and showed a predominantly axonal peripheral neuropathy, notably without renal or adrenal deficiencies ^19^.

Applying whole-exome sequencing to families with a complex neurological phenotype, we here identify a biallelic loss-of-function mutation in the *DEGS1* gene. Functional analyses in patient plasma and derived cells as well as in a chemical-and knockout model confirmed an increased dhSL/SL ratio and the formation of an atypical, novel SL metabolite as a result of DEGS1 insufficiency. Both findings are possible causes for the neurological symptoms in the patient. In summary, our study identifies a novel disease mechanism implicated in a human sphingolipid disorder that affects both the peripheral and the central nervous system.

## Results

### Clinical description and genetic analysis

The 22-year old male patient was first-born from healthy consanguineous Turkish parents and showed a progressive mixed pyramidal and extrapyramidal movement disorder as well as a progressive cerebellar atrophy. At the age of 6 months a motor developmental delay was observed and progressive spasticity developed in the further clinical course. Consecutive brain MRI revealed a general hypomyelination, a thinning of the brainstem and occipital white matter, severely reduced volume of both thalami, together with a progressive cerebellar and supra-and infratentorial atrophy, a thin corpus callosum, most pronounced in the dorsal part (**Fig. 1**). He developed a pathological EEG with epilepsy and grand mal seizures, which were successfully treated by a combination of valproate and carbamazepine. He showed a progressive neurological dysfunction, microcephaly, dystrophy, a progressive scoliosis, neurogenic bladder, and gastroesophageal reflux. Since age of 18 years feeding required a percutaneous endoscopic gastrostomy. Progressive spasticity resulted in flexion contractures of the extremities, a positive Babinski sign, and increased muscle tone. At the age of 19 years, an intrathecal baclofen pump therapy was initiated. Detailed clinical findings are summarized in **Table 1**. A muscle and N. suralis biopsy was performed at the age of 2 years. Archived electron micrographs (**Fig. 2**) from the sural nerve biopsy showed several nerve fibers with disproportionately thin myelin sheaths, moderate myelin folding, widening of the endoplasmic reticulum of Schwann cells and several autophagic vacuoles in the cytoplasm of Schwann cells. The muscle biopsy revealed neurogenic muscular atrophy according to the records that could be retrieved, however, no muscle specimens were available for review. Electroneurography at both arms and legs showed significant slowed nerve conduction velocities with only slight reduction of the amplitudes, in line with a predominant demyelinating neuropathy. Metabolic screening for lysosomal storage disorders did not show pathological findings. Genetic workup revealed a normal male karyotype (46, XY) and array-CGH was unsuspicious.

**Fig. 1:**
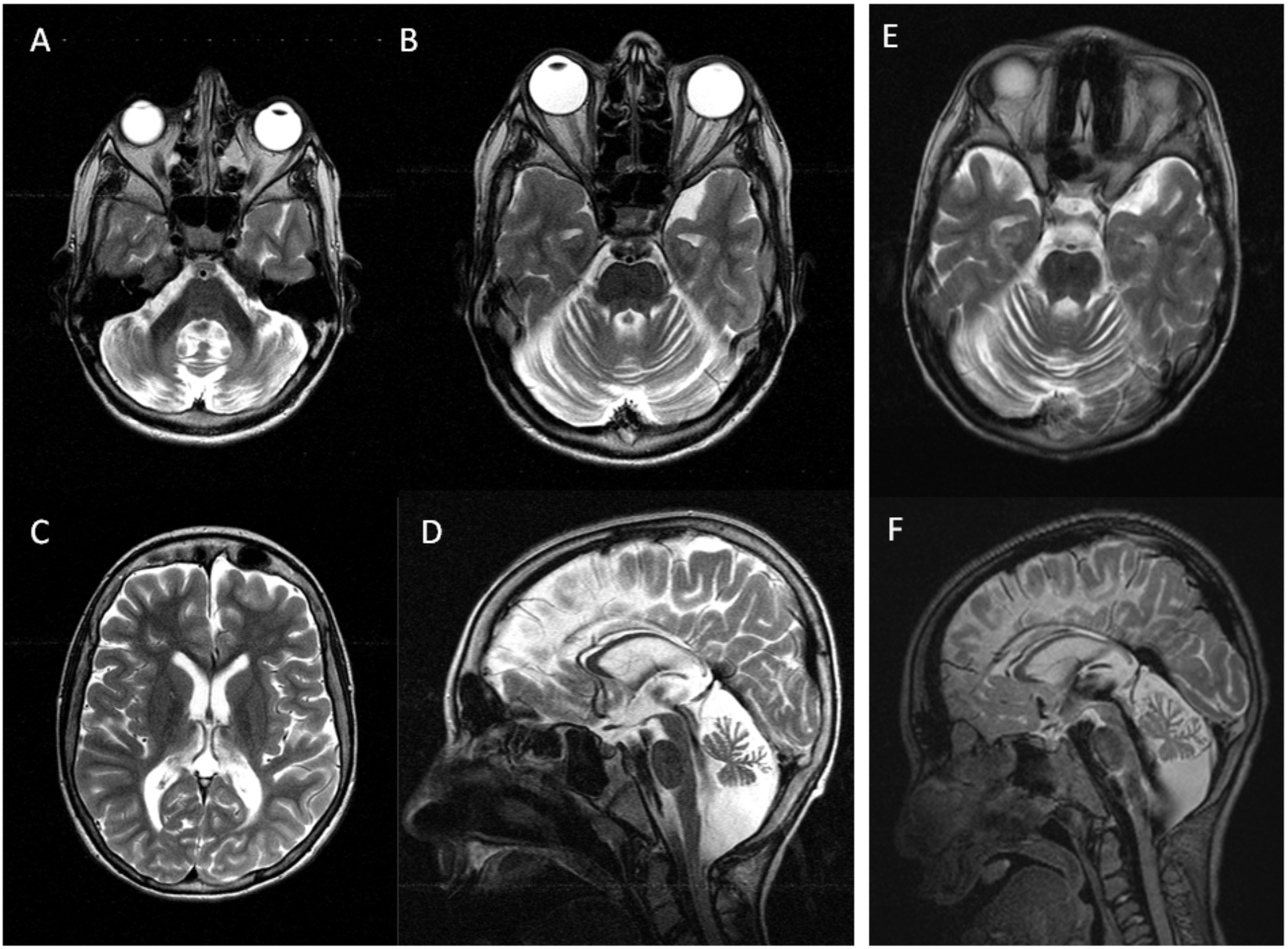
T2-weighted magnetic resonance imaging of the brain, axial (**A-C; E**) and sagittal (**D; F**), at 16 years of age (**A-D**) and 11 years (**E-F**). Severe and slowly progressive cerebellar atrophy with fiber degeneration of the middle cerebellar peduncles. The patient shows mild cortical atrophy and thin white matter especially in the posterior brain regions. In summary, MRI findings are in line with a progressive global neurodegenerative process.

**Tab. 1.** Clinical findings in the affected individual.

|  |  |
| --- | --- |
| DEGS1 (NM_003676.3) | c.839C>T; p.(Val280Ile) |
| Chromosomal position (hg19) | chr1:224380047C>T |
| Origin | Turkey (consanguineous parents) |
| Gender | Male |
| Age at onset | 6 months |
| Age at last follow-up | 21 years (y) |
| Clinical diagnosis | Multisystem disorder with progressive tetraspasticity, epilepsy, mental retardation, microcephaly |
| Clinical phenotype at last follow-up | Intellectual disability, non-ambulatory, severe spasticity, non-verbal, friendly |
| Body Weight | Normal at birth, dystrophy since the age of 3 y; at last follow-up 43 kg (-3.7 SD); percutan gastrostomy since age 18 y |
| Body Length | Normal at birth, mild short stature since the age of 4.5 y; at last follow-up 160 cm (-2.5 SD) |
| Head circumference | Normal in the first year than progressive microcephaly; at last follow-up 53 cm (-2.5 SD) |
| Epilepsy | Onset 5 years, grand mal epilepsy; under therapy with valproate and carbamazepin, since age 11 y no seizures |
| Brain MRI | Cerebellar atrophy, mild global atrophy thin white matter especially in the posterior brain regions |
| Visual evoked potentials | Pathological latency |
| Acoustic evoked potentials | Pathological latency |
| BERA | Pathological interpeak latency / auditory threshold 85 db |
| Electroneurography | Mild demyelinating neuropathy with decreased motoric nerve conduction velocities (mNCV): mNCV N. medianus 30.0 m/s age 2 y; 28.7 m/s age 12 y mNCV N. tibialis 38.0 m/s age 2 y; 39.2 m/s age 12 y |
| Nerve biopsy | Peripheral hypomyelination |
| Muscular biopsy | Neurogenic muscular atrophy |
| Scoliosis | Scoliosis surgery at age 18 y |
| Neurogenic bladder disorder |  |

**Fig. 2:**
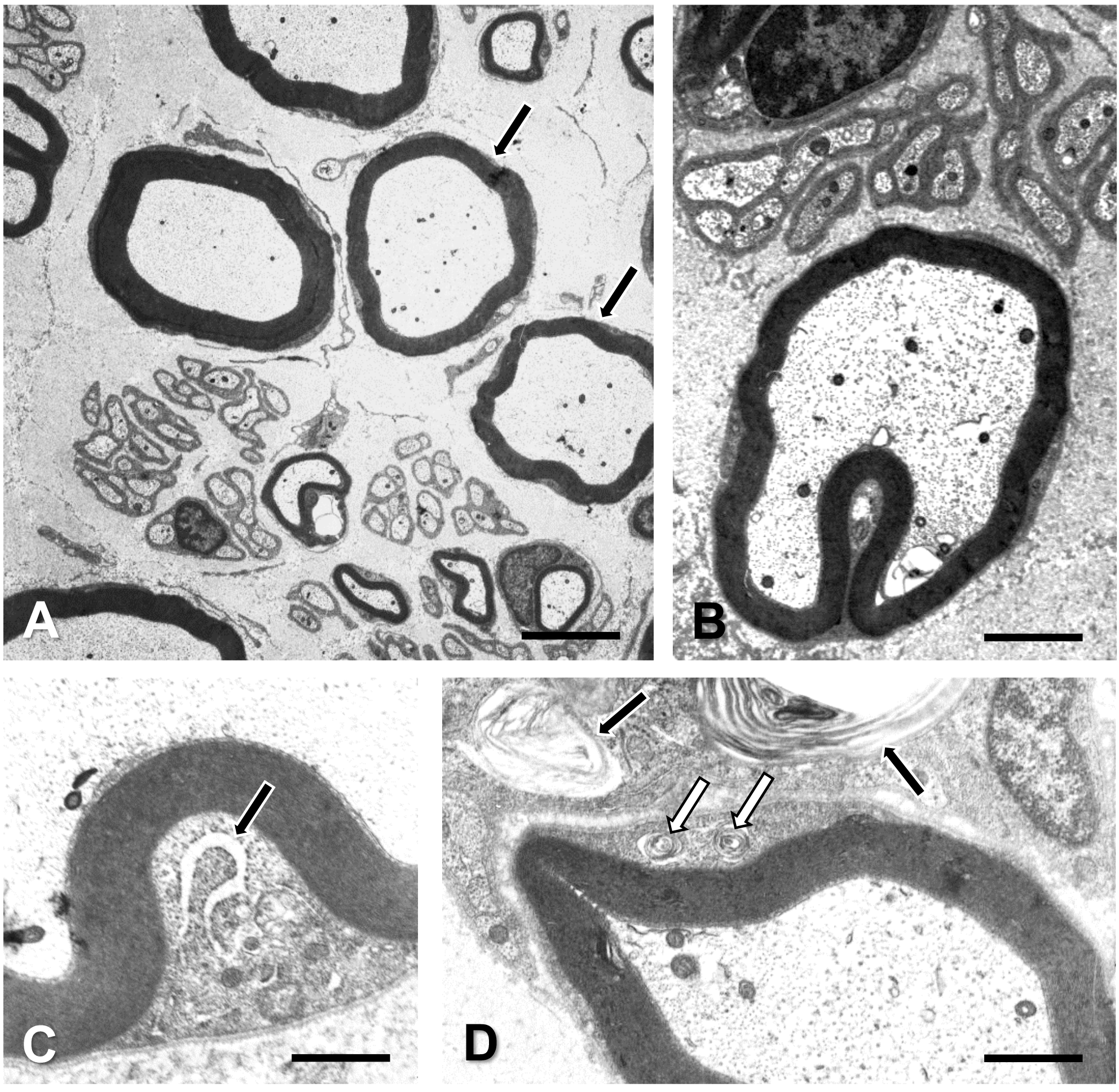
Electron micrographs of the sural nerve biopsy performed at the age of two years. **(A)** Nerve fibers with disproportionately thin myelin sheaths (arrows). Scale bar = 3 μm. **(B)** Occasional, moderate myelin folding. Scale bar = 1.8 μm. **(C)** Widening of the endoplasmic reticulum (arrow) of a Schwann cell. Scale bar = 0.5 μm **( D)** Small autophagic vacuoles in the cytoplasm of the Schwann cell of a myelinated nerve fiber (white arrows); black arrows: large autophagic vacuoles containing membranous debris in an adjacent cell which is covered by a basal lamina and may therefore be either a Schwann cell or a macrophage that has invaded a Schwann cell basal lamina sheath. Scale bar = 0.75 μm.

Using whole-exome sequencing in the index patient, his two unaffected siblings as well as both parents revealed a suspicious homozygous missense variant in the index patient in *DEGS1* (NM_003676.3) (**Fig.3** and **Suppl. Table 1**). Both parents and the siblings were heterozygous carriers of this *DEGS1* variant (**Fig. 3**). The variant changes the codon 280 from Alanine to Valine, p.(Ala280Val), affecting a highly conserved nucleotide (c.893C>T) and amino acid, which is located in the fatty acid desaturase / sphingolipid delta4-desaturase domain. The variant is not known to public databases (dbSNP, 1000genomes, ESP server, ExAC and GnomAD, June 2018) and is predicted to be deleterious by several bioinformatics pathogenicity prediction tools (CADD_phred (35), SIFT (0), Polyphen2_HDIV (0,99), MutationTaster prediction (D)).

**Fig. 3:**
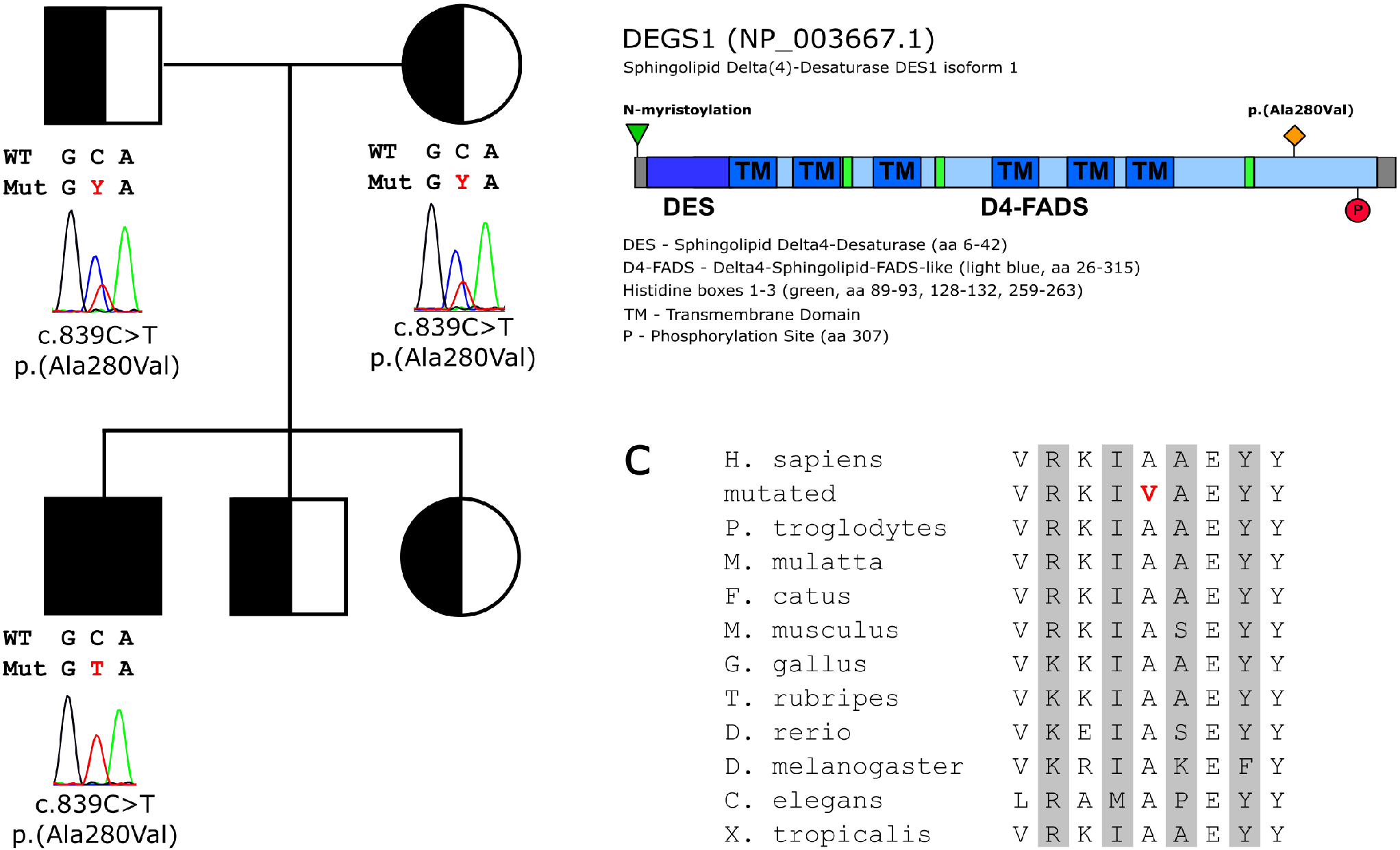
(**A**) The pedigree of the family shows the segregation of the *DEGS1* variant (NM_003676.3:c.839C>T, p.(Ala280Val), Chr1(hg19):g.224380047C>T) in the family. Sanger traces of the affected codon are shown in the index patient and his parents. (**B**) Domain architecture of the human DEGS1 protein. Position of the mutation indicated in orange. (**C**) Species alignment of the amino acid residues in proximity of the DEGS1 mutation. Mutation highlighted in red.

### DEGS1 subcellular localization

To investigate whether the p.(Ala280Val) missense-variant influences subcellular localization of DEGS1, the respective EGFP-tagged wildtype (wt) and variant p.(Ala280Val) proteins were heterologously overexpressed in HeLa cells and co-stained with antibodies specifically labelling the endoplasmic reticulum (ER) and mitochondria (**Fig. 4**). As expected, DEGS1 wt mainly colocalized with the ER marker protein disulfide isomerase (PDI) (Fig. 5A). No obvious difference regarding ER localization or ER morphology were observed upon expression of the mutant DEGS1 protein (**Fig. 4B**). Both, DEGS1 wt and p.(Ala280Val) showed little colocalization with the mitochondrial marker Tim23 (**Fig. 4C, D**).

**Fig. 4:**
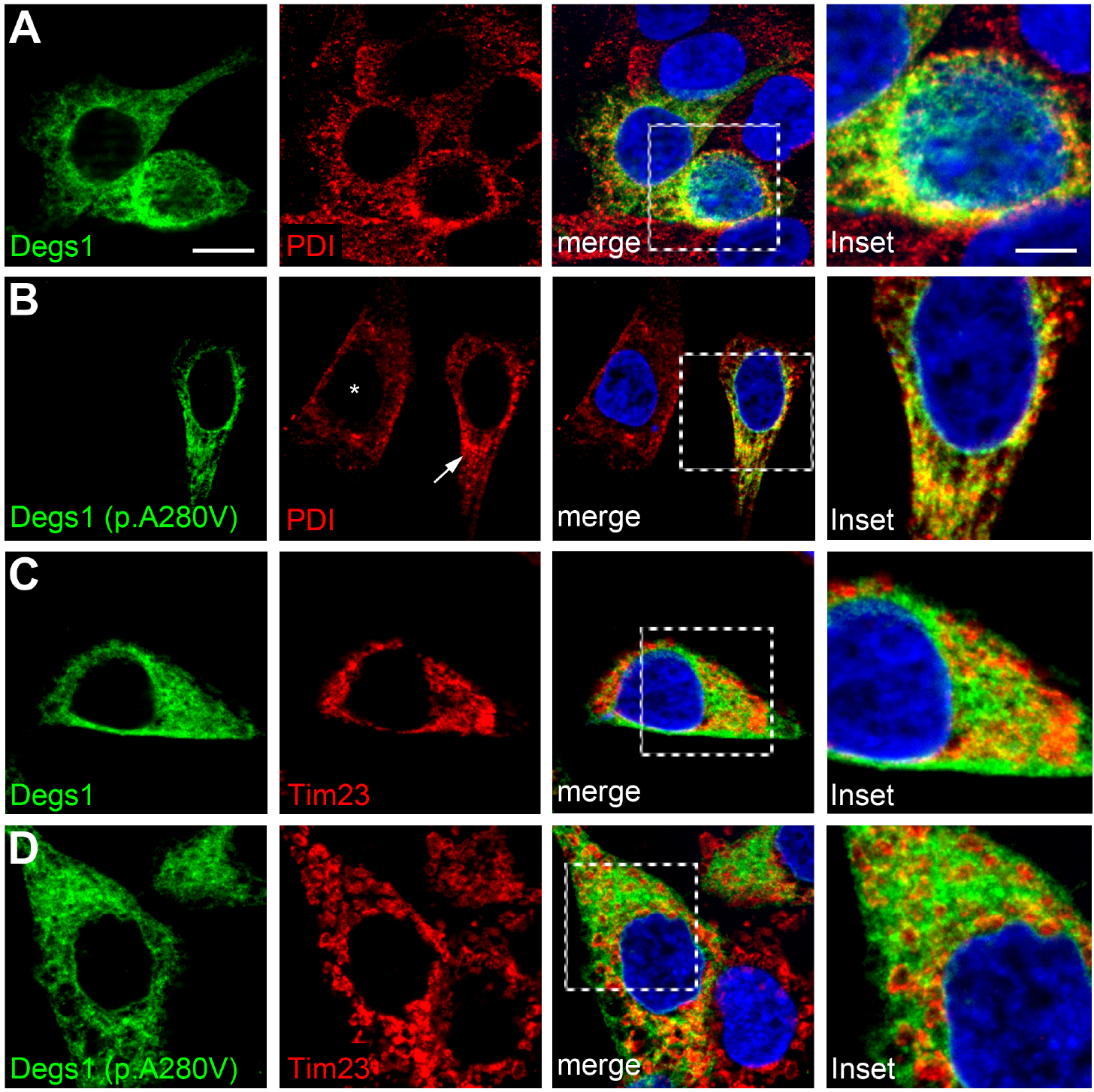
Cellular distribution of WT DEGS1 and the DEGS1 mutant p.(Ala280Val). (**A**) EGFP-tagged WT and (**B**) mutant DEGS1 colocalize with the endoplasmic reticulum-marker protein disulfide-isomerase (PDI). The reticular staining pattern of PDI in untransfected cells (asterisk in B) seems not disturbed in p.(Ala280Val)-overexpressing cells (arrow in B). (**C**, **D**) Overlap of immunofluorescence signals is observed for WT DEGS1 and the DEGS1 mutant p.(Ala280Val) overexpressing cells with the mitochondrion inner membrane marker Tim23. Scale bars: 10μm. Blow-ups 5μm scale.

**Fig. 5:**
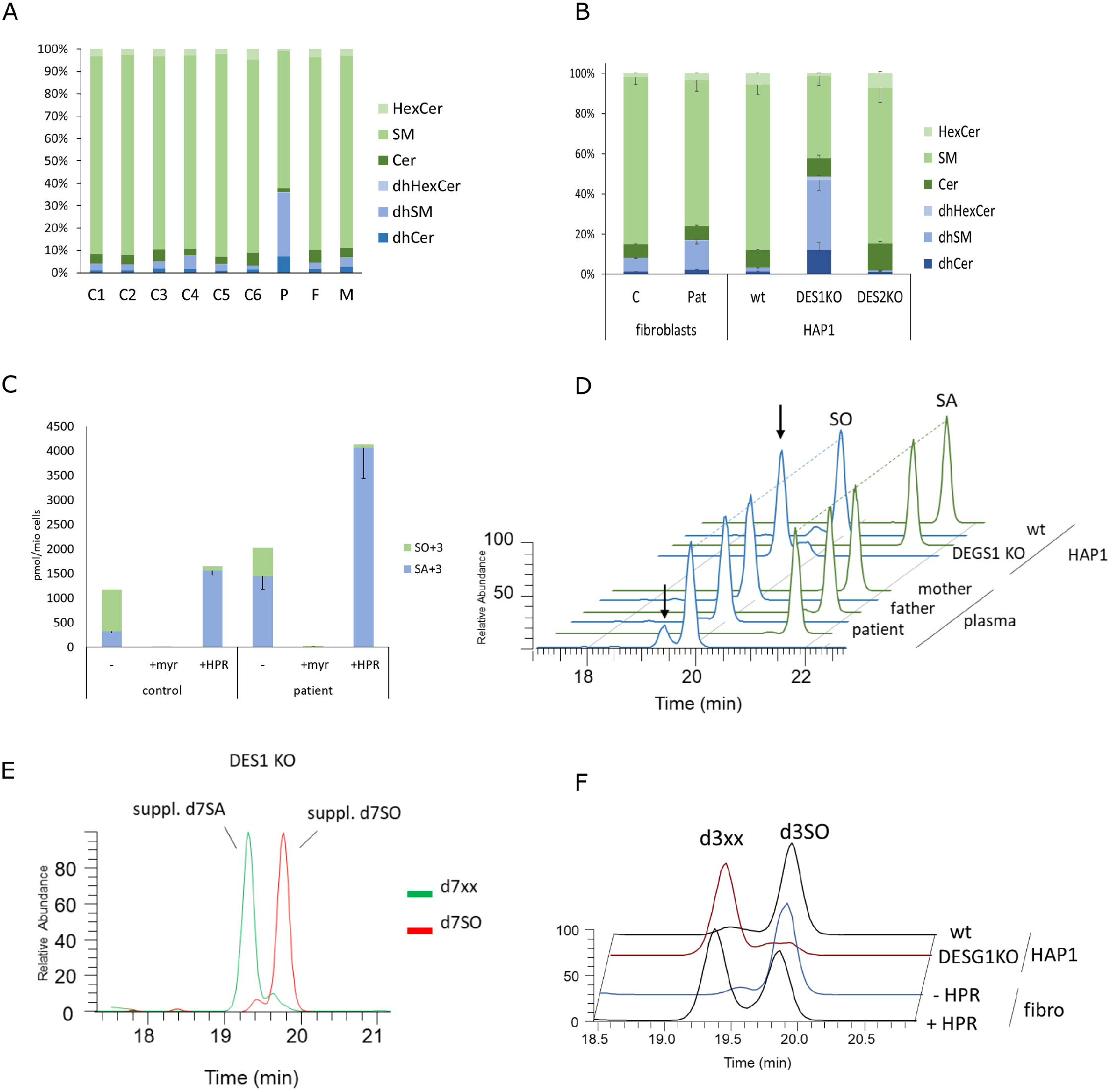
(**A**) Lipidomics proofing showed significantly elevated levels of dhSL species (dhCer, dhSM and dhHexCer) in patient plasma (P) compared to parents (F,M) or unrelated controls (C1-6). (**B**) Cultured patent derived fibroblast showed increased dhSL levels compared to cells from unrelated controls. Increased dhSL were also seen in HAP1 DEGS1KO cells where dhSL species made up to 50% of the total SL whereas the relative amount was less than 5% in HAP1 wt cells. No difference in the dhSL profile was seen in DEGS2 KO cells. (**C**) Isotope labeling assay of de-novo synthesized SLs. Control and patient fibroblast were supplemented with d4-serine (1mM) for 24h in absence or the presence of inhibitors for SPT (myriocin, myr) or DEGS1 (fenretinide, 4-HPR). The level of isotope labelled saturated species (SA+3) was significantly higher in the patent cells than in controls. No labelled sphingoid bases were formed in presence of myriocin (myr). When blocking DEGS1 activity with 4-HPR only the saturated (SA+3) species were formed. In the mutant cells and in presence of 4-HPR, total SL de-novo synthesis appeared to be increased.
(**D**) Identification of a novel, atypical sphingoid base peak in hydrolyzed patient plasma. The sphingoid base profile showed the presence of a novel sphingoid base (arrow) in patient plasma, which was not seen in the profile of the parents or unrelated controls. The metabolite was isobaric to SO (blue) but eluted about 30 seconds prior than normal SO. No second peak was seen for the saturated form (SA, green). The same metabolite was detected as peak in the DEGS1 KO but not in wt cells. (**E**) Characterization of the new metabolite as a downstream product of SA. DES1 KO cells were incubated for 24h with the isotope labelled sphingoid bases d7SA (1uM) and d7SO (1uM). The new peak appeared in isotope labelled form in d7SA - but not in d7SO treated cells. Besides, no conversion of d7SA into d7SO was observed in the DEGS1 KO cells indicating a complete loss of DEGS1 activity in these cells. (**F**) Isotopic labeling of the newly identified sphingoid base using d4-serine in fibroblast and in DEGS1 KO cells. In fibroblast, the labelled metabolite is only formed when DEGS1 activity was inhabited with 4- HPR. In d4-serine supplemented DEGS1KO but not in wt cells, the same peak was found d3 labelled.

### Sphingolipid analysis

A lipidomics analysis from plasma of the index patient, parent and six unrelated controls revealed a significant increase of dhSL species (sum of dhSM, dhCer and dhHexCer) in the patient plasma reflecting about 40% of the total plasma sphingolipids (**Fig. 5A**). The relative levels of dhSL species in the parent and controls were about 10% and varied only slightly between the individuals. A similar pattern, although less pronounced, was seen in cultured patient-derived fibroblast. At steady state levels, the dhSL species, in particular dhCer and dhSM, were significantly elevated in patient cells compared to skin fibroblast of unrelated controls (**Fig. 5B**). However, the increase in dhSL species in the patient fibroblasts was not as pronounced as in the plasma samples. To verify that this metabolic shift is caused by DEGS1 dysfunction, we used HAP1-knockout models (DEGS1 and DEGS2-deficient cells) generated by CRISPR/Cas technology. The lack of DEGS1 expression was confirmed by western blot (**Suppl. Fig. 1**). In the DEGS1-KO cells, about 50% of the sphingolipids were present in the dihydro (saturated) form. Despite this shift in the profile, the cells were viable and did not show differences in growth. In comparison, DEGS2-KO cells did not show an increase in dhSL species, indicating that the contribution of DEGS2 in the conversion of dhCer to Cer is minor (**Fig. 5C**). To estimate whether de-novo synthesis of sphingolipids is generally affected in the mutant cells, we performed an activity assay by measuring the incorporation of stable isotope labeled d4-serine over time. The added d4-serine is metabolized by SPT and incorporated into newly the synthesized sphingolipids. To facilitate the analysis and quantification of the labeled products, we performed a chemical hydrolysis of the extracted lipid prior MS analysis (see Methods). After hydrolysis, all saturated SL species are converted to SA+3 and all desaturated to SO+3. In control fibroblasts, the label was mostly found in the sphingosine form (**Fig. 5C**), whereas for patient fibroblasts the dominantly formed sphingoid base was sphinganine (SA+3). No labelled sphingoid bases were formed in the presence of the SPT inhibitor myriocin. However, in the patient cells about 25% of the labeled sphingoid bases were still converted to the desaturated form (SO+3) indicating some residual activity of the mutant. The remaining activity was further confirmed by chemical inhibition of DEGS1 with fenretinide (4-HPR). In presence of 4-HPR, >95% of the de-novo formed sphingoid bases were found in the dihydro form. In patient fibroblasts, 4-HPR led to a further reduction in SO+3 levels, which suggest that the p.(Ala280Val) mutant bears some residual enzyme activity which was further blocked with 4-HPR.

SPT activity and total SL de-novo synthesis appeared to be increased in the mutant cells and was further stimulated in presence of 4-HPR which suggest a dysfunctional metabolic feedback control of the pathway, when DEGS1 activity is reduced. However, total plasma levels were not elevated in the patient compared to parents or controls (data not shown). Surprisingly, when analyzing the sphingoid base profile in plasma after hydrolysis we identified a novel and atypical sphingoid base peak present in the patient plasma but not in the plasma of the parents or non-related controls (**Fig. 5D**). This additional metabolite had the same exact mass as SO but eluted about 30s earlier from the column. For SA we did not see an additional isobaric peak indicating that the new metabolite is formed downstream of SA. The same metabolite was formed in DEGS1-KO cells but was absent in HAP1 wt cells. Surprisingly, the peak was not visible in patient derived fibroblasts. Subsequent analysis showed that this novel metabolite is a bona-fide sphingoid base and formed as a downstream product of SPT. In d4-serine supplemented DEGS1-KO cells, this new metabolite was found to be d3 isotope labelled (**Fig. 5E**) and was absent in cells cultured in the presence of the SPT inhibitor myriocin (data not shown). The same isotopically labeled peak was also formed in control fibroblast in presence of 4-HPR (**Fig. 5E**), indicating that the presence of this metabolite is directly associated with a reduced DEGS1 activity. Interestingly, supplementing DEGS1-KO cells with either isotpe labelled d7SA or d7SO showed that this novel metabolite was specifically formed from d7SA but not from d7SO (**Fig. 5F**).

### Discussion

Here we identified *DEGS1* as novel gene implicated in a heritable multisystem disease with hypomyeliation and a degeneration of both the central and the peripheral nervous system. Early-onset developmental delay, movement disorder, progressive spasticity and epilepsy as disease hallmarks may be considered in mitochondrial disorders as well, but lactate values and muscle biopsy did not reveal changes typically for mitochondriopathies. Moreover, other multisystem neurological disorders such as neuronal ceroid lipofuscinosis, lysosomal storage disorders or leukodystrophies should be considered in the differential diagnosis of the here identified DEGS1-associated disease. We contacted neuromuscular centers in Belgium, the United Kingdom, and the U.S. to identify additional patients with *DEGS1*-mutations. However, re-evaluating available whole-exome data could yet not identify additional patients. Nevertheless, we suggest considering a DEGS1-associated disease in early onset hypomyelination, cerebellar atrophy and peripheral neuropathy and expect additional patients to be identified soon.

DEGS1 is a dihydroceramide desaturase and responsible for the conversion of dihydroceramide (dhCer) into ceramide by adding a Δ4,5 trans double bond in the sphingoid base backbone ^20^. DEGS2, a homologue isoform to DEGS1, was reported to act as a bifunctional enzyme (**Suppl. Fig 2**), both as a delta(4)-desaturase similar to DEGS1 and as a C4-monooxygenase adding a hydroxyl group to C4 position thereby forming phytosphingolipids^21^.

DEGS1 is a transmembrane protein residing in the endoplasmic reticulum (ER) membrane ^22, 23, 24^ that contains three conserved histidine-based motifs characteristic for membrane lipid desaturases and membrane hydrocarbon hydroxylases. Both, pharmacological and genetic ablation of DEGS1 activity critically impaired the ceramide synthesis pathway causing an increase in saturated, dihydro-sphingolipids (dhSL). As expected, in plasma of the index patient we found significantly elevated levels of dhCer, dhHexCer and dhSM species compared to the healthy parents or unrelated controls. A similar increase in dhSL species was seen in patient-derived fibroblasts and, independently, in a CRISPR/Cas-based *DEGS1* knockout HAP1 cell model as well as by chemical inhibition of DEGS1 with Fenretidine (4-HPR). The similar metabolic profiles in DEGS1-deficient cells indicate that the DEGS1 missense-variant has a reduced activity and is disease causing because of functional loss. However, comparing DEGS1 activity in patient fibroblast with DEGS1-KO HAP1 cells or in the presence of the DEGS1 inhibitor Fenretinide (4-HPR) indicates some remaining residual activity of the mutant. In mice, the homozygous deletion of DEGS1 (*Des1*^−/−^ mice) revealed an incompletely penetrant lethality. Surviving pups were small with a complex phenotype and multiple abnormalities including scaly skin and sparse hair, tremor, and metabolic abnormalities ^25^. *Des1*^−/−^ mice) had less ceramide and dramatically more dhCer in blood and tissues than wildtype littermates. However, heterozygous *Des1*^+/−^ animals were born at the expected Mendelian ratio and demonstrated no obvious health abnormalities. Interestingly, heterozygous *Des1*^+/−^ mice showed enhanced insulin sensitivity, normal glucose tolerance and were resistant to dexamethasone induced insulin resistance ^25^. This and other observations indicate that an increased dhCer/Cer ratio is linked to an improved glucose metabolism although the underling physiological mechanisms are not yet fully understood. Ceramides have long been considered as proapoptotic factors ^1^ which was demonstrated in a multitude of *in vitro* and *in vivo* models. However, the role of dhCer in apoptosis remained elusive up to now. Like ceramide, dhCer is not only found in the plasma membrane but also in other organelles such as ER, nucleus and mitochondria ^26^. Increased dhCer/Cer ratios result in increased rigidity of the plasma membrane ^27^, which likely affects many biological processes relying on appropriate membrane dynamics for active transport, vesicle formation, diffusion and signaling. It is tempting to speculate that the alterations in the ER structure as observed in the EM of the sural nerve of the patient may reflect impaired membrane properties. dhCer also prevents ceramide pore formation in planar membranes and mitochondria ^28^. However, defects in DEGS1 not only influence the dhCer/Cer ratio but also the profile of complex sphingolipids like (dh)SM and (dh)HexCer which are formed downstream of (dh)Cer. Sphingomyelin (SM) is an abundant component of glia and myelin and the different biophysical properties of dhSM might therefore significantly affect the physiology of glia cells and the structure of the myelin sheath which provides a direct link to the hypomyelination and neurological defects in the patient. In parallel to the increased dhSL/SL ratio, we also observed the appearance of a novel, atypical sphingolipid metabolite, which seems to be specifically formed under conditions of a reduced or absent DEGS1 activity. Lipidomics analysis showed that this metabolite is isobaric to canonical SO but eluted with a slightly shifted retention time compared to canonical SO. This isomeric peak was prominently seen in plasma of the index patient and in DEGS1-KO HAP1-cells as well as in 4-HPR treated cells. The newly formed metabolite becomes stable isotope labeled with d4-serine proving it as a bona-fide sphingolipid and downstream product of SPT and can also be directly formed from d7SA but not from d7SO. The chemical nature of this isobaric metabolite and its shift in the retention time suggests a different double bound (DB) position as canonical SO. Besides the typical Δ4-5E DB, some sphingolipids also contain a DB in Δ14-15Z position. This was first shown for SA-diene, a downstream metabolite of SO, and recently also demonstrated for 1-deoxySO, an aberrant sphingoid base formed by a mutant SPT enzyme in the context of sensory neuropathies (HSAN1) ^29^. We therefore hypothesize that this new metabolite could be a SO with a Δ14-15Z instead of the canonical Δ4-5E double bond. This metabolite might be formed in a metabolic bypass reaction from the accumulated dhCer by a not yet identified Δ14-15Z desaturase. The toxicity and pathophysiological relevance of this atypical SO isomer is not clear yet. However, it has been demonstrated that cis/trans change in the double bound position of ceramides significantly alters their biophysical and biological functions.

Assuming that the pathological and neurological defects in the patient are primarily caused by an altered dhSL/SL ratio, increasing the dietary consumption of animal sphingolipids might correct for this imbalance. In animals, sphingolipids are primarily found in the Δ4-5E form, which is reduced in the patient and found abundantly present in meat, milk and egg products. In contrast, plants and yeast usually have phytosphingolipids which bear a C4 hydroxyl group. An increased dietary consumption of SL from animal products might therefore increase the levels of unsaturated sphingolipid species and thereby normalize the dhSL/SL ratio in the patient. However, whether sphingolipid levels can be efficiently influenced by a dietary intervention and whether such a treatment also results in clinical improvement and/or a lack of disease progression remains to be tested.

In summary, we reported a novel human disease which is caused by a defect in the sphingolipid *de-novo* synthesis due to a mutant DEGS1 enzyme, leading to an altered dhSL/SL ratio and the formation of an atypical, aberrant and potentially toxic SL product.

## Methods

### Patient data

Written informed consent was obtained from the study participants after approval from the Institutional Review Boards at the participating institutions (Uniklinik RWTH Aachen: EK302-16). Consent was obtained according to the Declaration of Helsinki.

### Neuropathology

Resin embedding of the glutaraldehyde-fixed sural nerve biopsy tissue and subsequent electron microscopy had been performed using standard procedures ^30^.

### Whole-exome sequencing

Whole-exome Sequencing was performed with the DNA from peripheral blood of five family members including the index case, the two unaffected siblings and both parents. Enrichment was done with an Illumina Enrichment Kit (Nextera Rapid Capture Exome v1.2) and the respective libraries were sequenced on a NextSeq500 sequencer (Illumina, San Diego, USA). Alignment and variant calling was performed with SeqMule (v1.2) ^31^, (FastQC (version: 0.11.2), BWA-MEM (version: 0.7.8-r455), SAMtools (rmdup; version: 0.1.19-44428cd), SAMtools (filter; version: 0.1.19-44428cd), SAMtools (index; version: 0.1.19-44428cd, and GATKLite (realign; version: 2.3-9-gdcdccbb). Genome version hg19 was used for the alignment. Three variant caller were applied for variant detection (GATKLite UnifiedGenotyper (variant; version: 2.3-9-gdcdccbb), SAMtools (mpileup; version: 0.1.19-44428cd), FreeBayes (version: 0.9.14-14-gb00b735)). Variants called by at least two programs were considered for further analysis. The resulting variant files were combined (GATK, v3.6, CombineVariants) and processed with KGGSeq (v1.0, 14/Apr./2017) ^32^. Variants with a Minor Allele Frequency in public databases (i.e. ExAC, GnomAD, ESP, 1kG) above 0.75% were excluded. Average coverage in the target region was between 90-142× with 88-92% above 20 × coverage, respectively. Mutations were confirmed by sanger sequencing with BigDye Terminator 3.1 cycle sequencing kit and Genetic Analyzer 3500 (ThermoFisher Scientific, Darmstadt, Germany).

### Cells and cell culture

DEGS1 and DEGS2 knock out cells (DEGS1-KO HAP1, DEGS2-KO HAP1) were generated by a commercial service (Horizon Discovery, Waterbeach, United Kingdom). The introduction of a frame-shift mutation was confirmed by sequencing. HAP1 wt cells were provided as controls. HAP1 cells were cultured in Iscove’s Modified Dulbecco’s Medium (IMDM, ThermoFisher Scientific, Darmstadt, Germany) supplemented with 5 % fetal calf serum and 1 % penicillin/streptomycin at 37°C in a 5 % CO_2_ atmosphere. Patient fibroblasts were derived from a tissue biopsy in the index patient. HeLa (ATCC CCL-2) cells were cultured in Dulbecco′s modified Eagle′s medium (DMEM, ThermoFisher Scientific, Darmstadt, Germany) supplemented with 10 % fetal calf serum and 1 % penicillin/streptomycin at 37°C in a 5 % CO_2_ atmosphere.

### Immunohistochemistry

Transfection of HeLa cells (ATCC CCL-2) was done using Polyjet *in vitro* transfection reagent (SignaGen, Rockville, Maryland, USA) according to manufacturer′s protocol. Cells were fixed after 24 h with 4% paraformaldehyde in PBS. After blocking and permeabilization with 2% bovine serum albumine, 10% normal goat serum and 0.25 % TritonX-100 in PBS for 60 min at room temperature, cells were incubated with primary antibodies in blocking solution for 60 min at room temperature, washed 3 times in PBS and incubated with Alexa Fluor-568 or -647 secondary antibodies (1:1000; Molecular Probes). DAPI (4’,6-diamidino-2-phenylindole, 1:1000; Invitrogen) was used for nucleic acid staining. Images were taken with a Zeiss Observer Z.1 microscope equipped with an Apotome2 and HXP 120 lamp. The following primary antibodies were used: mouse-anti-PDI (Enzo, ADI-SPA-891, 1:200), mouse-anti-Tim23 (BD 611222, 1:200).

### Western blot

Protein isolation and Western Blot were carried out as described elsewhere^33^. The primary antibodies used for immunodetection were DEGS1 (Abcam, ab167169, 1: 5000) and α-tubulin (Abcam, ab15246, 1: 2000). As secondary antibody, horseradish peroxidase-conjugated antirabbit IgG (Santa Cruz, sc-2370, 1: 10000) was used. Detection was done using Clarity Western ECL Substrate (Bio-Rad, Munich, Germany) and FujiFilm LAS 3000 system (FujiFilm, Düsseldorf, Germany). PageRuler Plus Prestained Protein Ladder (ThermoFisher Scientific, Darmstadt, Germany) was used for protein weight estimation.

### Cloning

Human DEGS1 (NM_003676.3) was amplified from a commercially available cDNA library, using the following primers (Forward: caccatggggagccgcgtctcgcgggaagacttc, Reverse: ctccagcaccatctctcctttttggtg). The amplicon for DEGS1 lacking the stop codon was subcloned into pEGFP vectors (Clontech, Saint-Germain-en-Laye, France). The mutation p.Ala280Val was introduced by targeted mutagenesis. All inserts were sequence verified using Sanger sequencing. Primer sequences are available upon request.

### Lipidomics

Lipid extraction was performed as described previously ^34^ with some modifications. 20 μl plasma sample or 0.5-5 million cells were suspended in 20 μl PBS, 1 ml of a mixture of methanol: MTBE: chloroform (MMC) 4:3:3 (v/v/v) was added. The MMC mix was fortified with 100 pmoles/ml of the internal standards: d7-sphinganine (d18:0), d7-sphingosine (d18:1), dihydroceramide (d18:0:12:0), ceramide (d18:1/12:0, glucosylceramide (d18:1/8:0), sphingomyelin (18:1/12:0) and 50 pmoles/ml d7-sphingosine-1-phosphate. After brief vortexing, the samples were continuously mixed in a Thermomixer (Eppendorf) at 37 °C (1400 rpm, 20 min). Protein precipitation was obtained after centrifugation for 5 min, 16.000 g, 25 °C. The single-phase supernatant was collected, dried under N2 and stored at −20 °C until analysis. Before analysis, the dried lipids were dissolved in 100μl MeOH.

Liquid chromatography was done according to ^35^ with some modifications. The lipids were separated using a C30 Accucore LC column (Thermo Scientific, 150 mm * 2.1 mm * 2.6 μm) using the following mobile phases; A) Acetonitrile:Water (2:8) with 10 mM ammonium acetate and 0.1 % formic acid, B) Isopropanol: Acetonitrile (9:1) with 10 mM ammonium acetate and 0. 1 % formic acid and C) methanol at a flow rate of 0.3 ml/min.

The following gradient was applied; 0.0-1.5 min (isocratic 70 %A, 20 %B and 10 %C), 2. 1.5-18.5 min (ramp 20-100 % B), 3.18.5-25.5 min (isocratic 100 %B) 4. 25.5-30.5 minutes (isocratic 70 %A, 20 %B and 10 %C).

The liquid chromatography was coupled to a hybrid quadrupole-orbitrap mass spectrometer (Q-Exactive, Thermo Scientific) using a heated electrospray ionization (HESI) interface. The following parameters were used: spray voltage 3.5 kV, vaporizer temperature of 300 °C, sheath gas pressure 20 AU, aux gas 8 AU and capillary temperature of 320 °C. The detector was set to an MS2 method using a data dependent acquisition with top10 approach with stepped collision energy between 25 and 30. A 140000 resolution was used for the full spectrum and a 17500 for MS2. A dynamic exclusion filter was applied which will excludes fragmentation of the same ions for 20 sec. Identification criteria were 1) resolution with an accuracy of 5 ppm from the predicted mass t a resolving power of 140000 at 200 m/z. 2) Isotopic pattern fitting to expected isotopic distribution. 3) matching retention time and 4) the specific fragmentation patterns. Quantification was done using single point calibration. Pool samples in 4 concentration were used for quality control.

### Metabolic labelling and sphingoid base profiling

250.000 cells were seeded in 2 ml fresh medium in six-well plates (BD Falcon) and cultured for 2 days reaching ~70-80% confluence. The medium was exchanged for l-serine-and l-alanine-free DMEM (Genaxxon Bioscience, Ulm, Germany), containing 10% FCS, P/S. Two hours after medium exchange, isotope-labelled d3-N15-l-serine (1 mM) and (2,3,3,3)-d4-l-alanine (2 mM) was added (Cambridge Isotope Laboratories, Tewksbury, Massachusetts, United StatesI) In certain cases, Myriocyn (Focus Biomolecules • Plymouth Meeting, PA USA) or D7SA/D7SO (Avanti Polar Lipids, Alabaster, CA, USA) was also added to the cells. After 24 h, cells were harvested in 1 ml cold phosphate-buffered saline (PBS), counted (Beckman Coulter Z2), pelleted at 600 RCF at 4°C and stored at −20°C until further processing. During the SPT reaction, one deuterium is exchanged by a hydrogen, which results in newly formed sphingoid bases with a d3 isotope label. For the sphingoid base profiling the extracted sphingolipids were chemically hydrolyzed as described previously ^36^. During hydrolysis, the conjugated N-acyl chains and attached head groups are removed and the SL backbones released as free sphingoid bases. In consequence all saturated dhSL species are converted to SA and all unsaturated species to SO.

## Acknowledgements

We are grateful to the family participating in the study. The authors declare no conflict of interest. We thank Sebastian Gießelmann for excellent technical support. We thank Dr. Steuernagel, Klinikum Oldenburg, for initial fibroblast cultures. This work was supported by grants of the German Charcot-Marie-Tooth Disease Network (CMT-Net; 01GM1511D) to JW. Funding of the 7th Framework Program of the European Commission (“RESOLVE”, Project number 305707); the Swiss National Foundation SNF (Project 31003A_153390/1); the Hurka Foundation; the Novartis Foundation and the Rare Disease Initiative Zurich (“radiz”, Clinical Research Priority Program for Rare Diseases, University of Zurich) to TH.

## Author contributions

I. K., G.C.K. and T.H. designed the study. G.C.K., C.K., M.M. and M.E. assessed the phenotype of the patient. M. Bergmann, JMS and JW did the neuropathological analysis. Exome sequencing and evaluation was done by M. Begemann, F.K., I.K‥ Lipidomics studies were performed and evaluated by G.K., S.S., R.S. and T.H‥ Cell biological experiments were done by F.K., N.H. and B.H‥

## Additional information

### Supplementary Information

**Suppl. Table 1:** Whole-exome sequencing data

**Suppl. Fig. 1:** Western Blot (HAP1 wildtype and CRISPRS/Cas DEGS1-KO HAP1 cells)

**Suppl. Fig. 2:** Metabolic function of DEGS1 and DEGS2

## Competing financial interests

The authors declare no competing financial interests.

## Supplement

**Suppl. Fig. 1:**
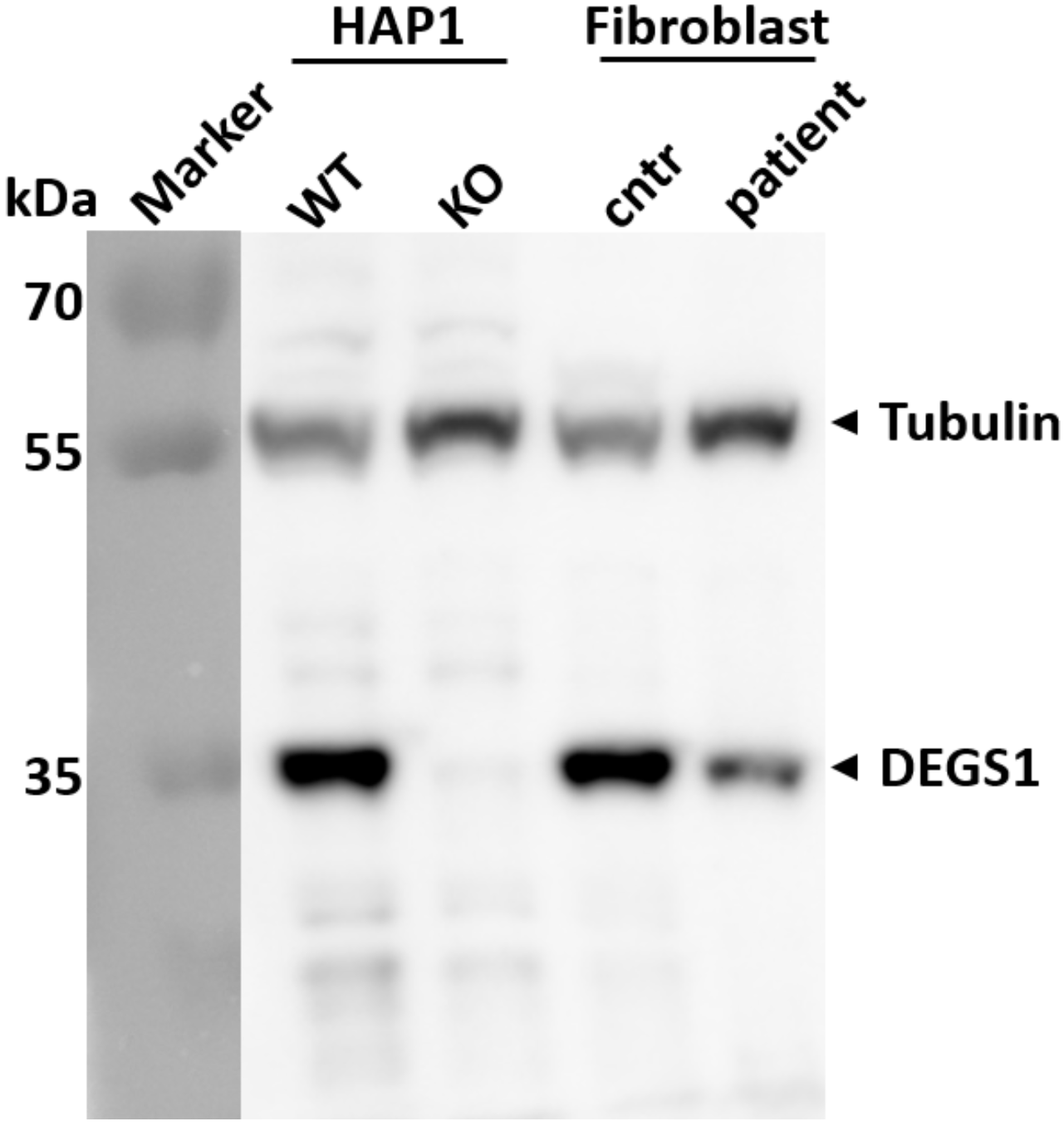
DEGS1 protein expression in HAP1 cell lines and patient fibroblasts. DEGS1 expression was analyzed by Western Blot in whole cell protein extracts from HAP1 cells (WT), DEGS1-KO HAP1 cells (KO), healthy control fibroblasts (cntr) and index patient fibroblasts (patient). Note absent DEGS1 protein in DEGS1-KO HAP1 cells and decrease in DEGS1 protein level in patient-derived fibroblasts in comparison to an unrelated healthy control, which may indicate an effect of the variant on protein expression or stability. kDa - kilo Dalton.

**Suppl. Fig. 2:**
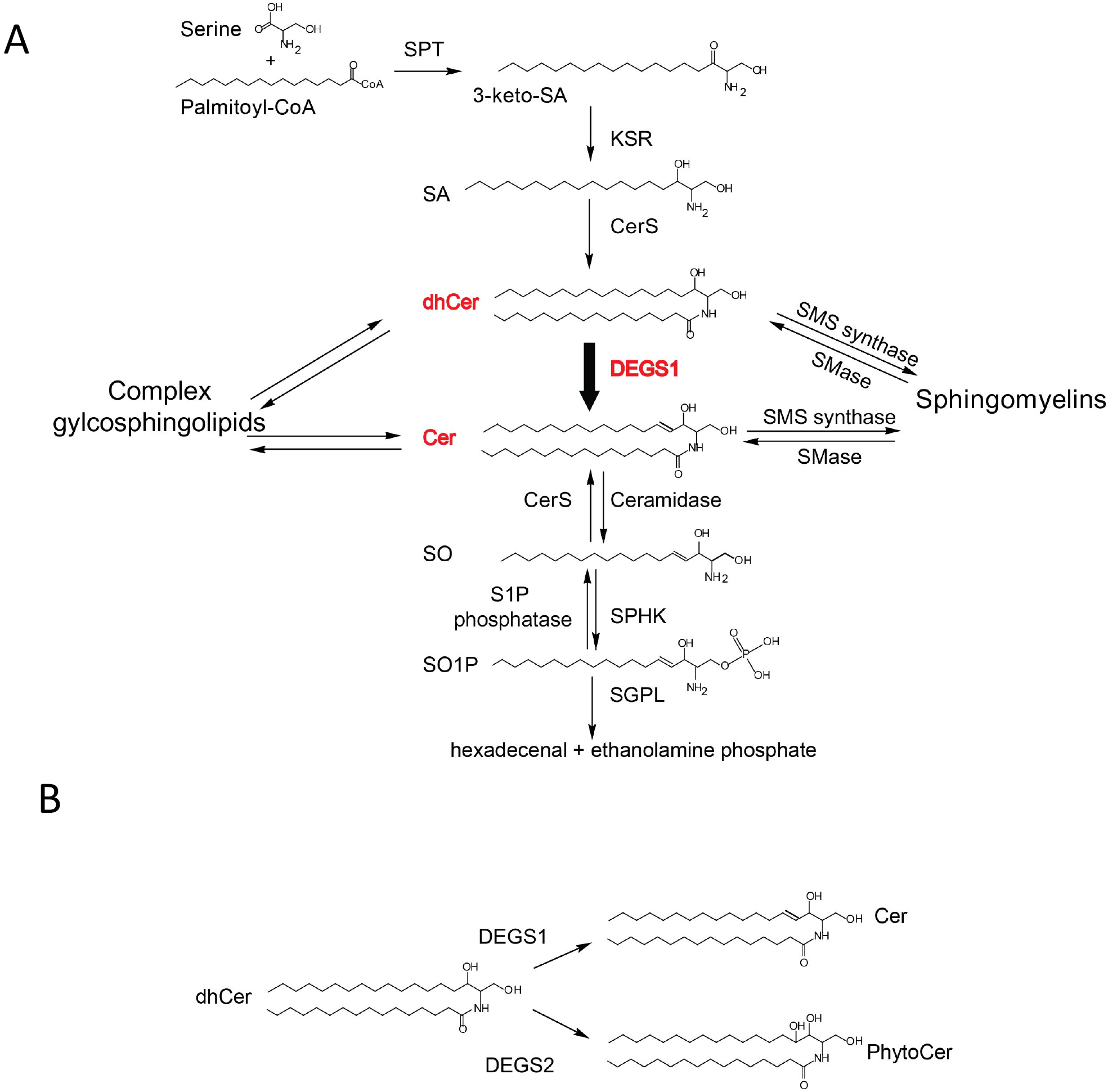
(**A**) Overview of the sphingolipid de-novo synthesis pathway. Complex sphingolipids such as SMs and GlucCer can be formed either from dh/Cer or Cer. (**B**) DEGS1 catalyzes the introduction of a Δ4-5E double bond into the sphingoid base backbone of dhCer thereby forming ceramide, whereas DEGS2 catalyzes the addition of an OH-group to the C4 position thereby forming phytosphingosine.

**Suppl. Tab. 1:** Whole exome data from five family members.

## References

1. Hannun YA, Obeid LM. Principles of bioactive lipid signalling: lessons from sphingolipids. Nat Rev Mol Cell Biol 9, 139–150 (2008).

2. Bikman BT, Summers SA. Ceramides as modulators of cellular and whole-body metabolism. J Clin Invest 121, 4222–4230 (2011).

3. Levy M, Futerman AH. Mammalian ceramide synthases. IUBMB Life 62, 347–356 (2010).

4. Coant N, Sakamoto W, Mao C, Hannun YA. Ceramidases, roles in sphingolipid metabolism and in health and disease. Adv Biol Regul, (2016).

5. Merrill AH, Jr., et al. Sphingolipids--the enigmatic lipid class: biochemistry, physiology, and pathophysiology. Toxicol Appl Pharmacol 142, 208–225 (1997).

6. Hanada K, et al. Molecular machinery for non-vesicular trafficking of ceramide. Nature 426, 803–809 (2003).

7. Bienias K, Fiedorowicz A, Sadowska A, Prokopiuk S, Car H. Regulation of sphingomyelin metabolism. Pharmacol Rep 68, 570–581 (2016).

8. Adada M, Luberto C, Canals D. Inhibitors of the sphingomyelin cycle: Sphingomyelin synthases and sphingomyelinases. Chem Phys Lipids 197, 45–59 (2016).

9. Sabourdy F, et al. Monogenic neurological disorders of sphingolipid metabolism. Biochim Biophys Acta 1851, 1040–1051 (2015).

10. Mosbech MB, et al. Reduced ceramide synthase 2 activity causes progressive myoclonic epilepsy. Ann Clin Transl Neurol 1, 88–98 (2014).

11. Vanni N, et al. Impairment of ceramide synthesis causes a novel progressive myoclonus epilepsy. Ann Neurol 76, 206–212 (2014).

12. Ferlazzo E, et al. Autosomal recessive progressive myoclonus epilepsy due to impaired ceramide synthesis. Epileptic Disord 18, 120–127 (2016).

13. Fragaki K, et al. Refractory epilepsy and mitochondrial dysfunction due to GM3 synthase deficiency. Eur J Hum Genet 21, 528–534 (2013).

14. Simpson MA, et al. Infantile-onset symptomatic epilepsy syndrome caused by a homozygous loss-of-function mutation of GM3 synthase. Nat Genet 36, 1225–1229 (2004).

15. Boukhris A, et al. Alteration of ganglioside biosynthesis responsible for complex hereditary spastic paraplegia. Am J Hum Genet 93, 118–123 (2013).

16. Janecke AR, et al. Deficiency of the sphingosine-1-phosphate lyase SGPL1 is associated with congenital nephrotic syndrome and congenital adrenal calcifications. Hum Mutat 38, 365–372 (2017).

17. Lovric S, et al. Mutations in sphingosine-1-phosphate lyase cause nephrosis with ichthyosis and adrenal insufficiency. J Clin Invest 127, 912–928 (2017).

18. Prasad R, et al. Sphingosine-1-phosphate lyase mutations cause primary adrenal insufficiency and steroid-resistant nephrotic syndrome. J Clin Invest 127, 942–953 (2017).

19. Atkinson D, et al. Sphingosine 1-phosphate lyase deficiency causes Charcot-Marie-Tooth neuropathy. Neurology 88, 533–542 (2017).

20. Causeret C, Geeraert L, Van der Hoeven G, Mannaerts GP, Van Veldhoven PP. Further characterization of rat dihydroceramide desaturase: tissue distribution, subcellular localization, and substrate specificity. Lipids 35, 1117–1125 (2000).

21. Mizutani Y, Kihara A, Igarashi Y. Identification of the human sphingolipid C4-hydroxylase, hDES2, and its up-regulation during keratinocyte differentiation. FEBS Lett 563, 93–97 (2004).

22. Cadena DL, Kurten RC, Gill GN. The product of the MLD gene is a member of the membrane fatty acid desaturase family: overexpression of MLD inhibits EGF receptor biosynthesis. Biochemistry 36, 6960–6967 (1997).

23. Sperling P, Zahringer U, Heinz E. A sphingolipid desaturase from higher plants. Identification of a new cytochrome b5 fusion protein. J Biol Chem 273, 28590–28596 (1998).

24. Ternes P, Franke S, Zahringer U, Sperling P, Heinz E. Identification and characterization of a sphingolipid delta 4-desaturase family. J Biol Chem 277, 25512–25518 (2002).

25. Holland WL, et al. Inhibition of ceramide synthesis ameliorates glucocorticoid-, saturated-fat-, and obesity-induced insulin resistance. Cell Metab 5, 167–179 (2007).

26. Andreyev AY, et al. Subcellular organelle lipidomics in TLR-4-activated macrophages. J Lipid Res 51, 2785–2797 (2010).

27. Vieira CR, et al. Dihydrosphingomyelin impairs HIV-1 infection by rigidifying liquid-ordered membrane domains. Chem Biol 17, 766–775 (2010).

28. Siskind LJ, Kolesnick RN, Colombini M. Ceramide forms channels in mitochondrial outer membranes at physiologically relevant concentrations. Mitochondrion 6, 118–125 (2006).

29. Steiner R, et al. Elucidating the chemical structure of native 1-deoxysphingosine. J Lipid Res 57, 1194–1203 (2016).

30. Nolte KW, Hans VJ, Schattenfroh C, Weis J, Schroder JM. Perineurial cells filled with collagen in ‘atypical’ Cogan’s syndrome. Acta Neuropathol 115, 589–596 (2008).

31. Guo Y, Ding X, Shen Y, Lyon GJ, Wang K. SeqMule: automated pipeline for analysis of human exome/genome sequencing data. Sci Rep 5, 14283 (2015).

32. Li MX, et al. Predicting mendelian disease-causing non-synonymous single nucleotide variants in exome sequencing studies. PLoS Genet 9, e1003143 (2013).

33. Esmaeili M, et al. The tumor suppressor ING1b is a novel corepressor for the androgen receptor and induces cellular senescence in prostate cancer cells. J Mol Cell Biol 8, 207–220 (2016).

34. Pellegrino RM, Di Veroli A, Valeri A, Goracci L, Cruciani G. LC/MS lipid profiling from human serum: a new method for global lipid extraction. Anal Bioanal Chem 406, 7937–7948 (2014).

35. Narvaez-Rivas M, Zhang Q. Comprehensive untargeted lipidomic analysis using coreshell C30 particle column and high field orbitrap mass spectrometer. J Chromatogr A 1440, 123–134 (2016).

36. Alecu I, et al. Cytotoxic 1-deoxysphingolipids are metabolized by a cytochrome P450-dependent pathway. J Lipid Res 58, 60–71 (2017).

